# Meniscal and ligament modifications in spontaneous and post-traumatic mouse models of osteoarthritis

**DOI:** 10.1101/816306

**Authors:** Lorenzo Ramos-Mucci, Behzad Javaheri, Rob van ‘t Hof, George Bou-Gharios, Andrew A Pitsillides, Eithne Comerford, Blandine Poulet

## Abstract

Osteoarthritis (OA) is a whole joint disease that affects all joint tissues, with changes in the articular cartilage (AC), subchondral bone and synovium. Pathologies in menisci and ligaments, however, are rarely analysed, although both are known to play vital roles in the mechanical stability of the joint. The aim of our study was to describe the pathological changes in menisci and ligament during disease development in murine spontaneous and post-traumatic surgically-induced OA and to quantify tissue mineralisation in the joint space using µCT imaging during OA progression.

Knees of Str/ort mice (spontaneous OA model; 26-40wks) and C57CBA F1 mice following destabilisation of medial meniscus (DMM) surgery (post-traumatic OA model; 8wks after DMM), were used to assess histological meniscal and ligament pathologies. Joint space mineralised tissue volume was quantified by µCT.

Meniscal pathological changes in Str/ort mouse knees were associated with articular cartilage lesion severity. These meniscal changes included ossification, hyperplasia, cell hypertrophy, collagen type II deposition and SOX9 expression in the fibrous region near the attachment to the knee joint capsule. Anterior cruciate ligaments exhibited extracellular matrix changes and chondrogenesis particularly at the tibial attachment site, and ossification was seen in collateral ligaments. Similar changes were confirmed in the post-traumatic DMM model. µCT analysis showed increased joint space mineralised tissue volume with OA progression in both the post-traumatic and spontaneous OA models.

Modifications in meniscal and ligament mineralisation and chondrogenesis are seen with overt AC degeneration in murine OA. Although the aetiology and the consequences of such changes remain unknown, they will influence stability and load transmission of the joint and may therefore contribute to OA progression. In addition, these changes may have important roles in movement restriction and pain, which represent major human clinical symptoms of OA. Description of such soft tissue changes, in addition to AC degradation, should be an important aspect of future studies in mouse models in order to furnish a more complete understanding of OA pathogenesis.

**Summary statement:** This manuscript describes histological changes in mouse knee joints in two models of osteoarthritis and correlates joint space mineralised tissue volume measured by µCT with disease severity.

## 1. Introduction

Osteoarthritis (OA) is a degenerative joint disease that leads to joint pain and restricted movement. Loss of articular cartilage (AC) characterises OA and is a hallmark of disease progression, but surrounding joint tissues including subchondral bone [1], synovium [2], menisci and ligaments are also affected [3]. Stability of the joint can influence OA progression and can be targeted to induce OA through surgical meniscectomy and ligament transection [4–6]. Despite the integral role of menisci and ligaments in maintaining normal joint function, few studies have examined their changing pathology during the development of disease in mouse models of OA [7–12]. Herein, we illustrate structural changes in the knee menisci and ligaments during OA development in two mouse models, including spontaneous (Str/ort mouse) and post-traumatic (destabilization of medial meniscus, DMM) OA models.

Though composition of menisci differs between mice and humans, menisci likely play similar stabilising roles in both species. Human knee menisci are fibrocartilaginous and contribute to load transmission and joint stability [13]. Unlike humans, mouse menisci are ossified and can be subdivided into three parts; a central bone core with marrow cavities, hyaline cartilage at the surface opposite tibial and femoral articulating surfaces, and an outer region of fibrocartilage tissue near the joint capsule [14]. In both species, the meniscus is attached to the femur and tibia by various ligaments [5].

Ligaments also stabilize knee joints by restricting excessive tibial movement relative to the femur in both animals and humans [15]. The cruciate ligaments prevent excessive antero-posterior and rotational movements, and the collateral ligaments restrict medio-lateral displacement [15]. Mouse ligaments have a similar structure and function to human ligaments [16]. The importance of ligament and meniscal stabilising functions is perhaps best exemplified by the induction of OA after ligament transection and meniscal injuries in a range of species, including both mouse [4, 5] and human [17–19]. Several findings also suggest meniscal [19] or ligament damage [20, 21] occurs in early stages of spontaneous OA. Despite these indications, meniscal and ligament pathology are not well understood.

Mouse models of OA are important tools to define mechanisms of disease and to determine targets to slow disease progression [22]. The main murine models for OA include spontaneous and mechanical induction models [23]. Many laboratory mouse strains develop OA with age [24]. The Str/ort mouse is known to exhibit early onset OA (visible OA seen by 18 weeks of age) with high incidence and severity, and reproduces many human OA features, such as proteoglycan loss, cartilage fibrillation, active degradation of cartilage extracellular matrix, osteophyte formation and subchondral bone thickening [8, 25–29]. These hallmarks are also well described in post-traumatic models of OA such as the DMM model, induced by surgical transection of the meniscotibial ligament [5, 30, 31]. Pathological changes in meniscus and ligaments are poorly described in these models of OA. Thus, the aim of this study was to explore the meniscal and ligament pathological changes during OA development in spontaneous and post-traumatic OA in mice, including tissue mineralisation in the joint space using micro-computed tomography (µCT) imaging and specific cellular and matrix markers of chondrogenesis using immunohistochemistry.

## 2. Results

### 2.1. Osteoarthritic meniscal changes in Str/ort mice include chondrogenesis and ossification of the outer region of the meniscus

To investigate the pathological changes in meniscus during osteoarthritis in mice, we used histological staining in healthy CBA mouse knee joint compared to osteoarthritic Str/ort mouse joints. Toluidine blue staining of the healthy CBA mouse knee joints (n=8) delineated the expected bone and hyaline cartilage surface components of the meniscus (Figure 1A, CBA). Additionally, Collagen type II (Col2; collagen specific to cartilage tissues) immunolabelling was present only in the hyaline cartilage surfaces of the meniscus, facing the articular femur and tibial cartilage. The chondrogenic marker SOX9 was sparsely expressed in the hyaline cartilage cells but not in the fibrous region.

**Figure 1:**
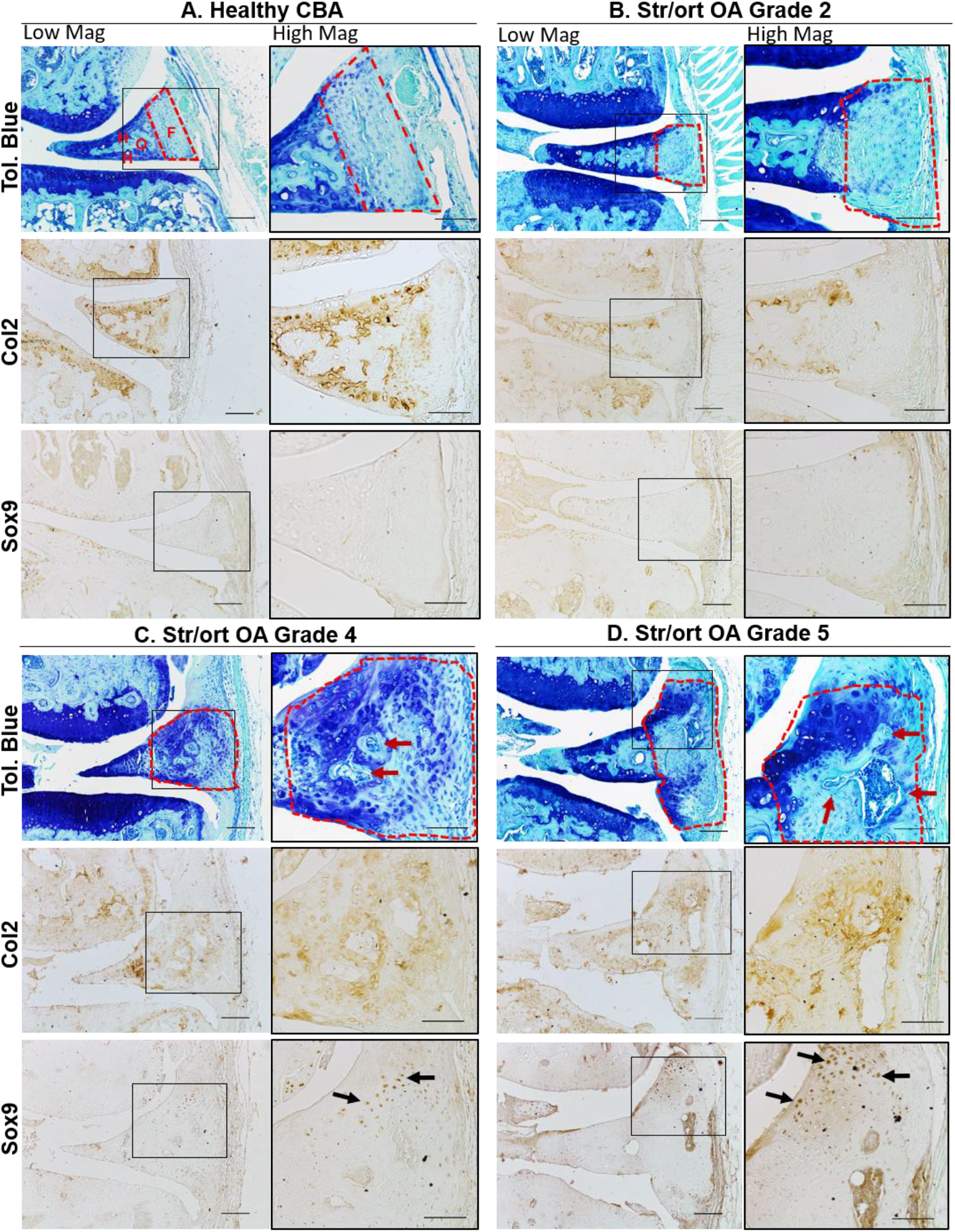
Representative images of meniscal pathology during spontaneous osteoarthritis development in Str/ort mice. A. Toluidine blue, Collagen type II (Col2) and Sox9 immunolabelling in healthy CBA mouse meniscus, which can be divided into distinct regions: hyaline cartilage (H) surrounds an ossified region (O) and an outer fibrous (F) region. B-D. Representative images from diseased menisci from Str/ort mouse knee joints with OA grades of 2 (mild), 4 (moderate) and 5 (severe). Toluidine Blue staining showed a range of meniscal pathologies associated with OA development including an increase in the fibrous region (delineated by red lines), proteoglycan deposition and bone formation (red arrows). Col2 and sox9 positive cells (black arrows) were seen in the fibrous region of the meniscus with disease showed. Low mag =low magnification, scale bar = 100µm; High mag = high magnification, scale bar = 50µm. For orientation: femur at top of picture, tibia bottom.

Knee joints from a group of male adult Str/ort mice (aged 26 and 40 weeks, n=21) were graded for AC degradation using the OARSI grading system showing a range of severities (grades 2-6). The range of meniscal pathology occurred mainly in the medial compartment and included osteophyte formation at the meniscal tip, chondrogenesis and ossification in the fibrous region, as well as hypertrophy at attachment site and erosion. The severity of pathological changes in the medial meniscus of Str/ort mice increased with AC degradation severity scored according to the OARSI grading system (Figure 1B-D). The first notable change in early stages of OA (grade 2) was an increase the outer fibrous region of the meniscus, which further increased with disease progression. In OA grade 4 and higher, this expanding fibrous region showed evidence of hyperplasia, chondrogenesis and formation of bone marrow cavities. The changes included increased toluidine blue staining and the appearance of rounded chondrocyte-like cells particularly in the region attaching to the joint capsule. Immunohistochemistry showed Col2 deposition in the fibrous region, particularly in areas surrounding bone formation and in the outer areas near the capsular attachment site. Similarly, Sox9 positive cells were prominent in the edges of the fibrous compartment of the meniscus.

### 2.2. Cruciate and collateral ligaments developed areas of chondrogenesis during osteoarthritis in Str/ort mouse knees

Ligaments, the stabilizers of the knee joint, are comprised of aligned spindle-shaped cells and dense collagenous fibres which stain only weakly with toluidine blue in healthy joints (Figure 2A). The anterior cruciate ligament (ACL) in Str/ort joints with AC lesions of grade 2 or greater, contained ACLs with clear chondrogenic changes including positive toluidine blue staining, thickening of fibres, and rounded hypertrophic cells evident at the tibial insertion (Figure 2). In more severe OA cases, positive toluidine blue staining expanded through the entire ACL and insertion site and showed signs of misalignment and degeneration, with columns of hypertrophic cells. Immunohistochemistry confirmed changes in the ACL extracellular matrix and cellular markers in the Str/ort mouse knee joint, with Col2 deposition in the ACL tibial insertion sites (Figure 2). SOX9 expression was seen within cells of the ligament insertion site and extended to the cells within the ligaments (Figure 2). Col2 deposition and SOX9 expression in the ligament or insertion site was not observed in the healthy CBA ligaments (Figure 2).

**Figure 2:**
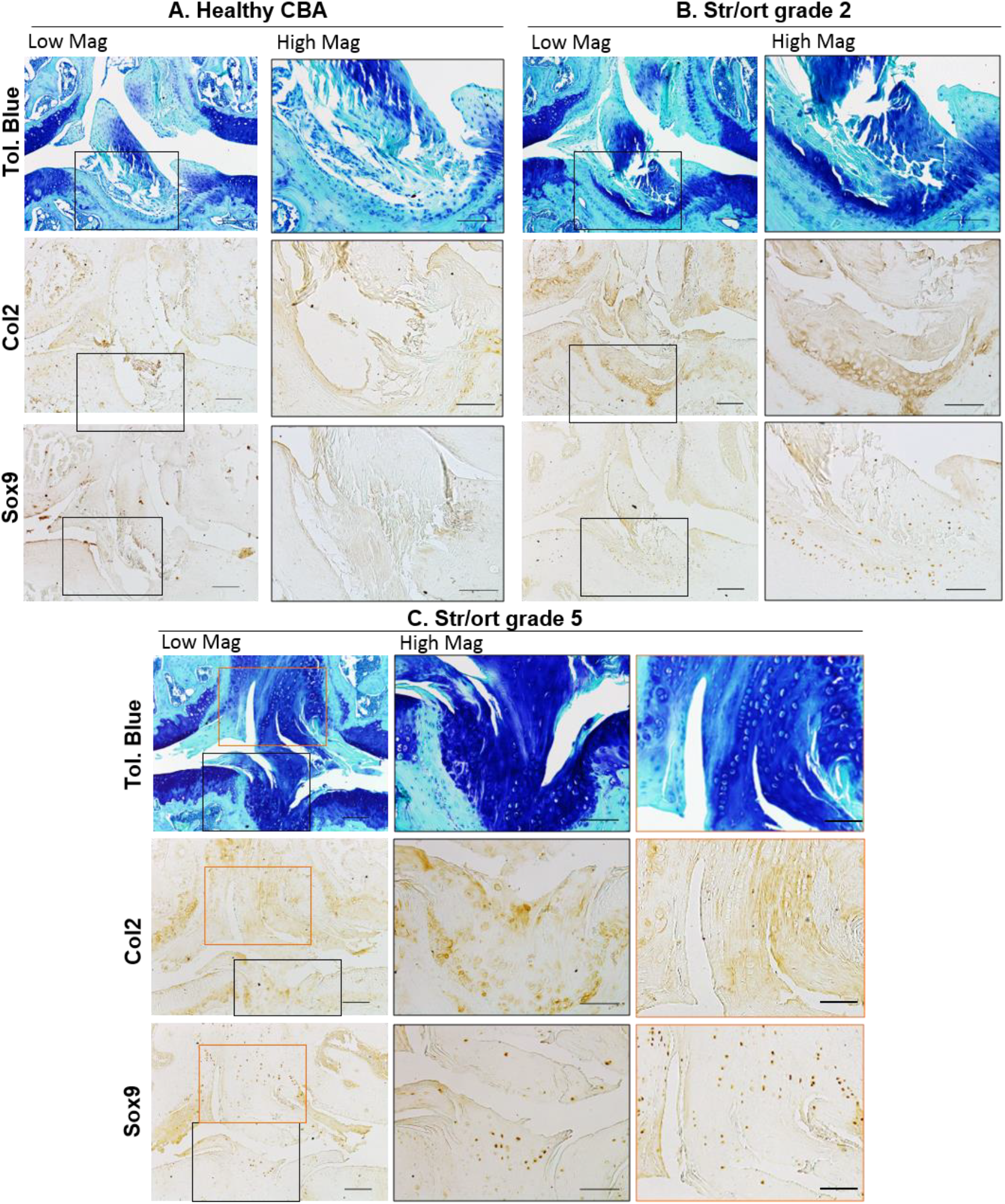
Representative images of cruciate ligament changes with OA development in Str/ort mice. A. Toluidine blue, Collagen type II (Col2) and Sox9 immunolabelling in healthy CBA mouse cruciate ligaments, with high magnification of insertion site into the tibia. B-C. Cruciate ligaments from Str/ort mouse knee joints with OA grades of 2 (mild) and 5 (severe). Toluidine Blue staining showed increased staining at the insertion site (High mag for panels B and C) and within the ligament body (High mag for panel C), with hypertrophy of local cells. Col2 deposition and sox-9 positive cells were also increased, especially in severe diseased joints. Low mag =low magnification, scale bar = 100µm; High mag = high magnification, scale bar = 50µm. For orientation: femur at top of picture, tibia bottom.

Collateral ligaments link the femur and tibia on the lateral/medial sides of the joints. Similarly to ACL, cells in collateral ligaments in healthy CBA joints are spindle shaped and form small columns along the ligament fibres, which are devoid of col2 deposition (Figure 3A). Cells within the body of the ligament have very mild SOX9 expression. In Str/ort mice, collateral ligaments showed signs of overt thickening, increased toluidine blue staining, cell hypertrophy and areas of bone formation (Figure 3). These features are consistently accompanied by positive local labelling for col2 deposition and sox9 expression. In very severely osteoarthritic joints, marked structural changes with significant bone nodule formation can be seen; in these joints, the attachment site is also modified with bone formation extending into the ligament (Figure 3C).

**Figure 3:**
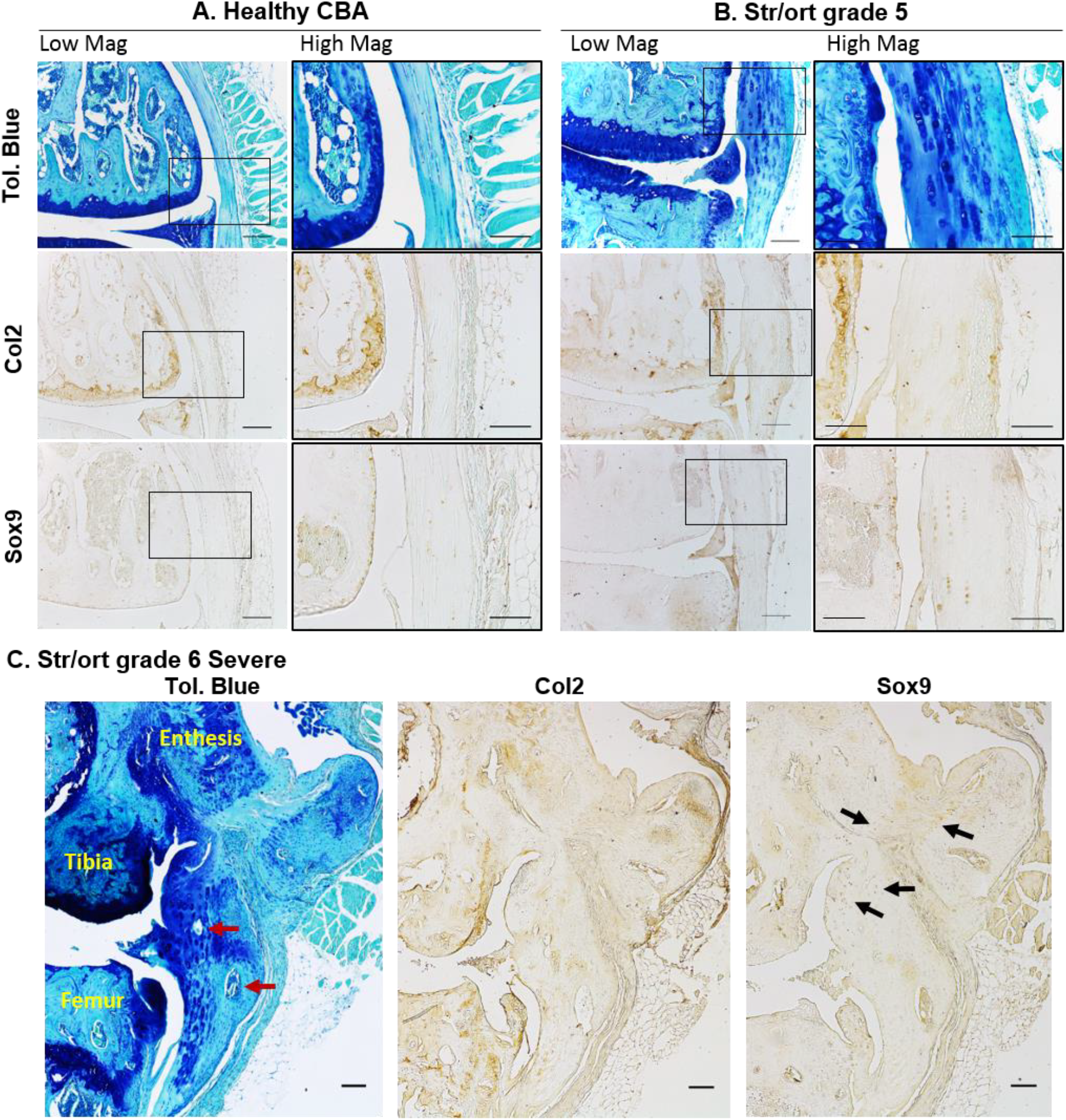
Representative images of collateral ligaments in OA Str/ort mouse knee joints. A. Toluidine blue, Collagen type II (Col2) and Sox9 immunolabelling in healthy CBA mouse collateral ligaments, with high magnification of the body of the ligament. B. Collateral ligaments from Str/ort mouse knee joints with OA grades of 5 (severe). Toluidine Blue staining showed increased staining with hypertrophy of local cells. Col2 deposition and sox-9 positive cells were also increased. C. Very severe OA joints showed severe changes including ossification within the body of the ligament (red arrows) and areas of col2 deposition and Sox9-expressing cells (black arrows). Low mag =low magnification, scale bar = 100µm; High mag = high magnification, scale bar = 50µm. For orientation: femur at top of picture, tibia bottom.

### 2.3. Histological analysis of a post-traumatic OA mouse model reveals chondrogenic and hyperplasic changes in the menisci and ligaments

The post-traumatic model of osteoarthritis (DMM) (n=13) lead to moderate OA development 8 weeks post-surgery, with an average AC degradation OARSI grade of 3.4±0.3, primarily in the medial compartment. Similar meniscal changes to Str/ort mouse knee joints were seen in this port-traumatic model of OA (see Table 1). The medial meniscus showed hyperplasia of meniscal attachments seen as a significant thickening, and chondrogenesis followed by ossification in the outer fibrous compartment, including blood vessel invasion and the formation of marrow spaces (Figure 4). Toluidine blue staining was also noted in the areas surrounding bone formation and near the capsular attachment along with evidence of cell hypertrophy. The tibial insertion site of the ACL displayed increased proteoglycan staining with toluidine blue and evidence of cell hypertrophy (Figure 4). The ACL also showed signs of disorganisation, including thickening and misalignment of the fibres.

**Table 1:** Summary of meniscal and ligament changes in two murine models of knee osteoarthritis. Maximum OA grades reported for Str/ort mice based on the OARSI grading system, to group these changes according to severity of OA in a model that shows variability.

|  | *Spontaneous OA*<br>*(Str/ort mouse)* | *Surgical model*<br>*(DMM)* |
| --- | --- | --- |
| ***Meniscus*** |  |  |
| *Outer part chondrogenesis* | Grade 3-4 | Yes |
| *Outer part ossification* | Grade 3-5 | Yes |
| *Hypertrophy of attachment* | Grade 4-6 | Yes |
| *Erosion* | Grade 6 | Yes (Grade 6) |
| ***Ligaments (cruciates, collaterals)*** |  |  |
| *Strong ECM staining* | Grade 3-6 | Yes |
| *Cell hypertrophy* | Grade 3-6 | Yes |
| *Loss of alignment* | Grade 4-6 | Yes |
| *Ossification* | Grade 4-6 | Yes |

**Figure 4:**
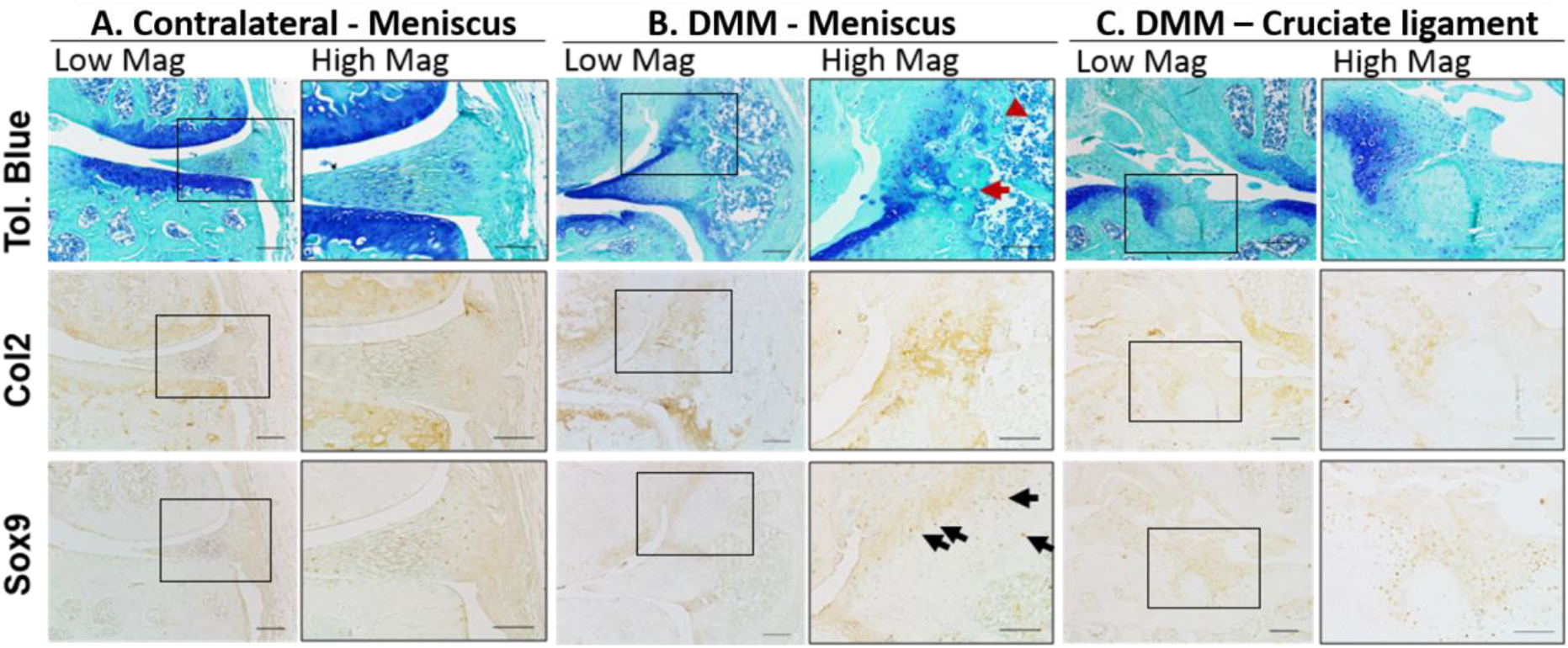
Representative images of meniscus and ligament modifications during post-traumatic OA in DMM joints. A. Staining of the contralateral control meniscus. B. Toluidine Blue staining showed meniscal changes in the DMM mice associated with OA development, including bone formation in the fibrous meniscal attachment site, proteoglycan and bone deposition (red arrow). Col2 deposition was also seen at the sites of high toluidine Blue staining with Sox9-expressing cells (black arrows). C. Cruciate ligament insertion site in DMM joints showed areas of strong toluidine blue staining, concomitant with col2 deposition and Sox9 expression. Low mag =low magnification, scale bar = 100µm; High mag = high magnification, scale bar = 50µm.

### 2.4. Quantification of joint space mineralisation by µCT analysis reveals significant increases in mineralisation volumes in osteoarthritic mouse knees

Mineralised tissues can be visualised in 3-dimensions (3D) and quantified using X-ray based µCT imaging techniques. Using this technique, we aimed to quantify the changes described above to the soft tissues, including ligament and meniscal cartilage/bone formation. To achieve this, we used knee joints (n=8) from Str/ort mice with a range of cartilage lesion severities (from grade 3 to 6), from C57CBA F1 mice at 8 weeks following DMM surgery (n=7) and contralateral healthy joints, for µCT analysis of the joint space. These analyses showed that Str/ort mouse knee joints developed increased mineralised tissue volume within the joint space with increasing severity of cartilage lesion grades in Str/ort mice (Figure 5A). Similarly, an increase in mineralised tissue volume was also found in response to DMM surgery compared to the untreated contralateral leg (Figure 5C).

**Figure 5:**
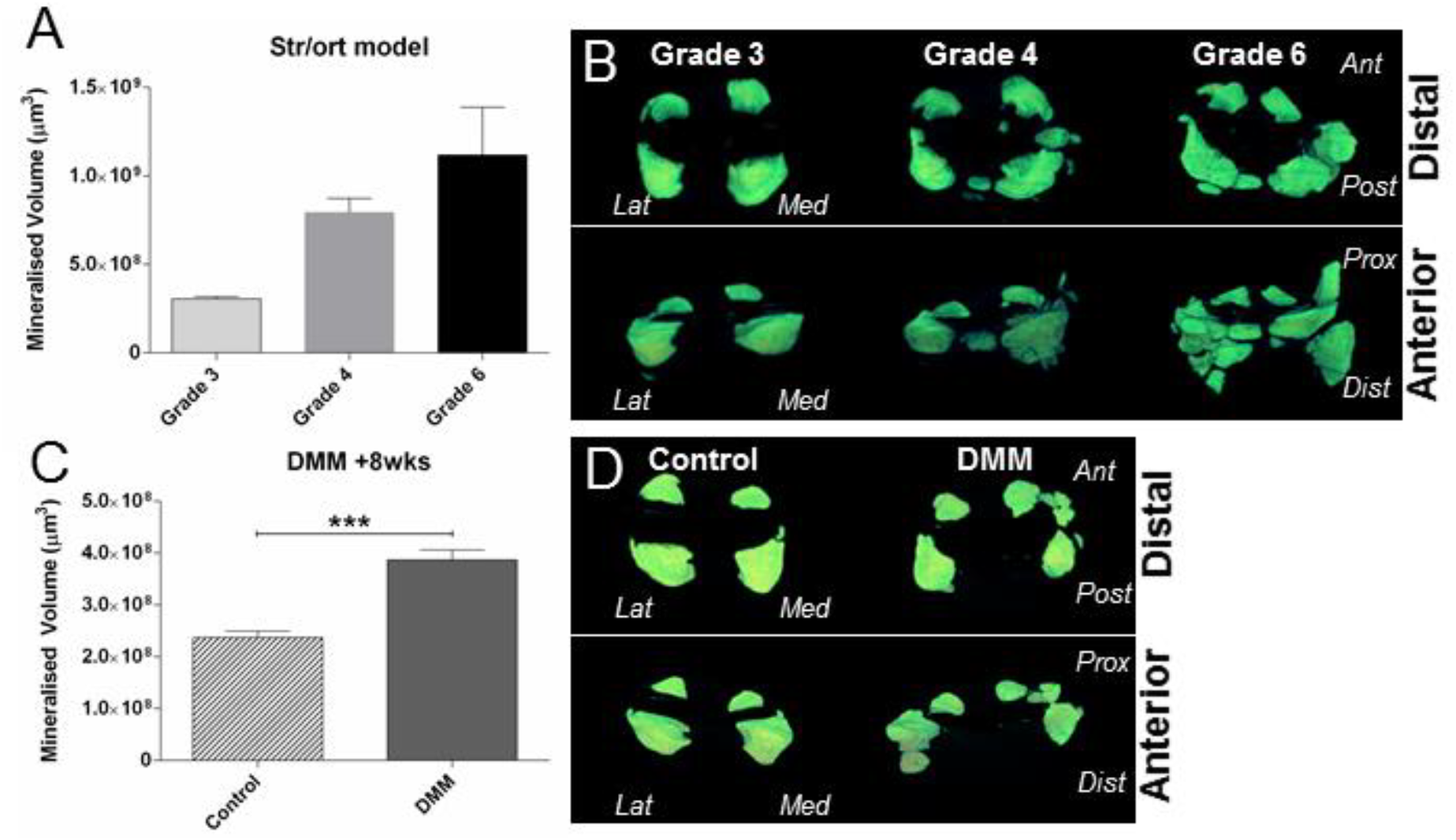
µCT-based quantification and 3D images of joint space mineralisation with OA progression in Str/ort mice and in response to post-traumatic DMM surgery. (A) Analysis showed volume of mineralised tissue in the joint space increased with disease severity in Str/ort mice. (B) This increase was also visualised in 3D as enlargement of the meniscus and ectopic mineralisation nodules of different sizes (Distal and Anterior views). (C) Similar increases in mineralised tissue volume was noted eight weeks following DMM surgery compared to the contralateral control knee, (D) with mineral nodules primarily in the medial side of the joint. (Lat = Lateral; Med= Medial; Ant = Anterior; Post = Posterior; Prox = Proximal; Dist = Distal). *** for p>0.001, based on t-test between Control and DMM. Data presented as mean ± SEM.

Representative 3D images of this mineralized tissue however showed differing patterns of pathological mineralisation between these models. The Str/ort mice joints showed early changes in the medial compartment, which is also the primary location of cartilage lesions, but developed to more widespread abnormal mineralisation in distinct nodules around both menisci in severe OA (Grade 6; Figure 5B). In contrast, the DMM model, which induces OA mainly in the medial compartment, showed increased mineralisation primarily around the medial meniscus compared to contralateral untreated control limbs (Figure 5D).

Overall, µCT analysis of the mouse knee showed quantifiable mineralisation changes in the joint space with OA progression.

## 3. Discussion

This study describes the pathological changes to the menisci and ligaments during osteoarthritis development in mouse models of OA, including spontaneous Str/ort mouse and surgical post-traumatic DMM models. We found abnormal patterns of chondrogenesis, ossification, cell hypertrophy, meniscal erosion and loss of fibre alignment in ligaments in both models. Pathological changes in both meniscal and ligamentous tissues indicated endochondral ossification markers included collagen type II matrix formation and SOX9 expression. Moreover, we described the use of µCT analysis to quantify these changes in soft tissue mineralisation/ossification with OA development.

Similar studies have reported changes in non-articular cartilage tissues during OA development in mice. Some of the meniscal changes reported herein have been briefly reported in Str/ort mice [7–9] and more recently in the DMM mouse model [32], with a general histological analysis but no description of the ACL. The description of knee osteoarthritis in Str/ort mice by Walton and others [8, 9] includes reports of heterotopic calcification. In particular, they describe an increased ossification centre in the medial meniscus [8] and a positive association between medial collateral ligament calcification and OA development [9]. These findings agree with our histological data of meniscal ectopic mineralisation and ossified nodules in the collateral ligaments in Str/ort mice. Anderson-MacKenzie et al showed that Str/ort mice ACLs have increased MMP-2, high collagen remodelling and decreased mechanical strength compared to healthy CBA joints in 27 week-old Str/ort mice, at which time histological OA is well underway [20]. Meanwhile, Kwok *et al* showed murine DMM menisci with severe fibrillations, erosion and calcifications [32]. This also matches our histological results and is further strengthened by our novel µCT mineralisation measurements and our immunohistochemistry, which provides quantification of these changes and insight into the matrix and cellular changes involved.

Col2 and SOX9 markers are typically cartilage-specific markers crucial for endochondral ossification processes. With the exception of ligament enthesis insertion sites [33], positive expression these cartilage markers are not expected in normal ligaments. Healthy menisci express these cartilage markers on the cartilage surface of the main body of the meniscus [11, 14], but not in the outer compartment of the meniscus, where most of the OA changes we describe occur. Similarly to our findings, Col2 and SOX9 have been associated with OA development in human ACLs [34] and confirmed the chondrogenic nature of our observed increased toluidine blue staining in the meniscus and ligament in murine OA.

In humans, meniscal pathologies have been associated with ageing and OA. Indeed, the meniscus commonly develops lesions in OA patients with no previous history of injury [35]. Although meniscal ossicles are rare in humans and arise mainly from mechanical trauma [36], calcification is more common and found mainly in OA patients [37, 38]. In addition, aged and OA menisci showed changes in cellularity (with both areas of hyper- and hypocellularity), cell clustering and phenotype changes from fibroblast-like cells to round chondrocyte-like cells [38], similar to our findings in mouse OA. Similarities in meniscal pathologies between human and mouse OA support a favourable translatability between the species, despite clear meniscal structural differences (i.e. ossified meniscus in mouse). Pathological meniscal ossification has also been described in other animal models of OA. In the guinea pig model of spontaneous OA, for example, the extent of meniscal ossification has been found to correlate significantly with disease severity [39]. In the same model, treatment with an anti-mineralization agent blocked meniscal calcification and cartilage damage [40]. Meniscal ossification is also an important feature in osteoarthritic dogs and pigs [41, 42]. In pigs, meniscal ossification was associated with abduction of the hindlegs, stiff locomotion and reduced weight gain, indicative of a painful condition [41]. On the other hand, in a rat model of DMM, histological scoring found no differences in meniscal ossification between sham and DMM groups [43]. Incidence of increased meniscal ossification in a majority OA species confirms the need to determine the mechanisms involved that could have broad implications for controlling OA development.

The pathological chondrogenesis of ligaments described in mouse knee joints in this study also shows many similarities to human joints [44, 45]. It has been found that human cruciate ligaments from OA patients show important chondroid and cartilage metaplasia, which involve a change in ligament cell phenotype to a more chondrocyte-like round cell morphologies [34, 46]. This included positive Col2 and ColX labelling in areas of chondroid metaplasia along with SOX9 and RUNX2 expression in chondrocyte-like cells [34]. The mechanisms of pathological chondrogenesis and ossification in these ligaments remain, however, largely undefined.

In addition to a link with OA development [47–51], factors linked to ligament ossification include Indian hedgehog, Wnt and inflammatory cytokine signalling [44, 45, 52]. It was also reported that COL6A1 and RUNX2 may be common susceptibility genes in ligament ossification in the spine [53, 54] and both of these genes have also been linked to OA [55, 56]. TGFβ and BMPs are known regulators of ossification and fibrosis as well as being involved in OA development, and have also been associated with ligament ossification, including recombinant BMP2 in rat spines [57] and abnormal BMP signalling in human spines [58]. Recently, epidermal growth factor signalling was linked to OA development, comprising of ectopic chondro-osseous pathologies in the ligament and meniscus [10]. Mechanical trauma or disturbances are possible factors that contribute to these pathological ossifications, with evidence that ossification of ligaments is increased in response to mechanical loading [59–62]. Our study describes changes in a model of spontaneous OA (Str/ort); although the aetiology of OA development in this model is largely unknown, it does not preclude the possible changes in the joint mechanical environment during disease development in this model that could promote these ligament pathologies. In fact, similarities in the meniscal and ligament pathogenesis between spontaneous and post-traumatic models of OA suggest a common pathway for these events. Future work in mouse models will define important pathways involved in these pathologies to target OA development.

It is still uncertain, however, whether such meniscal and ligament changes contribute to OA progression. They might influence joint stability and load transmission, therefore accelerating OA and restricting the range of movement. Furthermore, damage in ligaments and menisci in human elderly patients has been linked with knee pain [63], so the associated ossification, with neo-angiogenesis and innervation, may increase joint pain [64] possibly leading to gait modifications. Further studies in murine models of OA might help decipher the aetiology of pain and restricted movement - the main clinical manifestations in human OA, as well as providing a better understanding of OA as a whole joint organ disease.

Most meniscal and ligament pathologies in OA are generally based on histological description, which are qualitative and at best semi-quantitative based on a scoring system of specific histological hallmarks [65, 66]. In this study, we have used µCT imaging to better quantify changes in these tissues. We found that there was a clear increase in mineralised tissue volume in the joint space and OA progression in each model, based on either AC lesions scores or disease progression following time after induction. The reproducibility of the increases in mineralised tissue volume with OA in both models suggests that this method may be a strong predictor of disease severity and progression. This new method could also be used as a non-invasive marker of disease progression in mice *in vivo*.

Our description of meniscal and ligament tissue modifications with the advancement of OA in two different mouse models suggests that these changes are an important hallmark of disease. The reproducibility of these events and the possible functional implications in these innervated tissues, such as mechanical disturbances and pain, emphasizes the need for further studies and for systematic reporting of these events in future *in vivo* studies. In addition, changes in joint space mineralised tissue volume measurements during OA development, suggest an important new method of following and quantifying disease progression using non-invasive µCT imaging techniques.

## Materials and Methods

### Animals

Male CBA (Charles River, UK), Str/ort (in-house, Royal Veterinary College, London, UK) and C57-CBA F1 mice (Charles River, UK) were kept in polypropylene cages, subjected to 12hour light/dark cycles, at 21±2°C and fed standard RM1 maintenance diet ad libitum (No.1; Special Diet Services, Witham UK). All procedures complied with Animals (Scientific Procedures) Act 1986 and local ethics committee.

Male Str/ort mice aged 26 (n=7) and 40 weeks (n=14) were used; this age range allowed the full extent of OA development encompassing a range of severity grades of disease (from grades 2 to 6). Thus, changes in the meniscal and ligament tissues were assessed based on grade of AC lesion severity as a measure of OA severity using the internationally-recognised OARSI grading system [65]. For the µCT analysis, a second group of Str/ort mice were used (n=8) all at the same age in order to minimise the effect of age on meniscal size. These mice were used at 37 weeks of age, which showed variability in OA severity grades ranging from grades 3 to 6 [67]. For toluidine blue and immunohistochemical staining, Str/ort samples were compared to aged-matched CBA controls (n=6 per group).

### DMM

Surgical destabilisation of the medial meniscus was performed on the right knee joint in n=13 male C57CBA F1 mice at 10 weeks-old as described before [68]. Briefly, anaesthesia was induced by injection of 10 μl/g of Hypnorm®/Hypnovel® (at a ratio of 1:1–4 parts water), the right knee joint was accessed via a medial incision. The meniscotibial ligament was transected, resulting in the release of the medial meniscus from its tibial attachment. After transection, the skin was sutured and mice were immediately transferred to a heated post-operative recovery room. All animals received buprenorphine HCl (Vetergesic; Alstoe Animal Health, York, UK) sub-cutaneously post-surgery and monitored daily to ensure good health. For µCT analysis, toluidine blue and immunohistochemical staining samples were compared to contralateral controls.

### µCT

cadaveric knee joints were scanned with a 5µm isotropic voxel size (50kV, 200µA respectively, 0.5mm Aluminium filter; 0.6° rotation angle, no frame averaging) using a Skyscan 1172 µCT scanner (Skyscan, Belgium). Hand-drawn regions of interests of the whole menisci (lateral and medial and other mineralised tissues that were not part of the tibial or femoral bones were analysed using 3D algorithms in CTAn (Skyscan, Belgium) to provide the mineralised tissue volume (measured as Bone Volume on CTAn). Statistical analysis of mineralised volume comparing the different groups within each OA model was performed using a student t-test for the DMM, and ANOVA with Bonferroni post-hoc test for the Str/ort mouse model. Three-dimensional models of the menisci were created using CTVox from the region of interest selected for mineralised tissue volume analysis (Skyscan, Belgium).

### Histology

Animals were killed by cervical dislocation and knee joints prepared for µCT imaging and/or histology. Briefly, skin and muscles were removed, the joints fixed in neutral buffered formalin, stored in 70% ethanol and scanned as described above. For histology, joints were decalcified (ImmunocalTM, Quarttet, Berlin, Germany), dehydrated and processed for wax embedding. Serial coronal 6µm thick sections were cut across the entire joint and a quarter of the entire set from regular intervals across the joint stained with toluidine blue (0.1% in 0.1M solution of acetate buffer, pH 5.6) and counterstained with 0.2% fast green for 5 seconds. Toluidine blue was used for histological examination to assess pathophysiological changes in menisci and ligaments.

### Immunohistochemistry

Immunohistochemistry was performed to localise expression of collagen type II (Col2) (Thermo, Mouse MC) and SOX9 (Abcam, Rabbit PC). Histology slides were dewaxed and rehydrated. For Col2, antigen retrieval was applied with pepsin (3mg/mL in 0.02M HCl) for 45 min at 37°C. Slides were then washed and blocked for endogenous peroxidase with 0.3% hydrogen peroxide for 15 min at 37°C (Sigma). Next, slides were blocked for endogenous Avidin/Biotin binding with an Aviding/Biotin Blocking Kit (Vector Labs, SP2001). Non-specific binding sites were blocked for 1 hour (Col2: MoM Kit, Vector Labs, BMK-2202; Sox9: 10% v/v goat serum). Primary antibodies were incubated overnight at 4°C and included Col2 (1/100, Thermo) and Sox9 (1/1000, EMD Millipore). Negative controls included a mouse IgG (2µg/mL, Sigma, for Col2) or rabbit IgG (1µg/mL, Vector Labs, for SOX9). Following washing, biotinylated secondary antibody (Vector Labs) was applied for 1 hour and then Vectastain (Vector Labs) for 30 min. Stains were developed with DAB (Vector Labs), dehydrated and mounted with DPX (Sigma).

## Author contribution

Study concept and design: B.P., A.P.

Acquisition, analysis and interpretation of data: all authors. Statistical analysis: B.P, L.R.

Drafting of the manuscript: all authors.

Critical revision of the manuscript: all authors.

